# Fluctuations in AAV genome quantification via digital PCR affected by heat-based virion lysis

**DOI:** 10.1101/2025.09.04.674220

**Authors:** Antonina Govorov, Filip Trajkovski, Nazguel Wagner, Lucie Klughertz, Marine Guise, Karl Pflanz, Alexandra Müller-Scholz, Robert Hertel

**Affiliations:** Sartorius Lab Instruments GmbH & Co. KG, Otto-Brenner-Straße 20, 37079 Göttingen, Germany; Sartorius Stedim Biotech GmbH, Otto-Brenner-Straße 20 37079 Goettingen, Germany; Sartorius Polyplus SAS, 75 Rue Marguerite Perey 67400 Illkirch, France; Department for Genomic and Applied Microbiology, Institute of Microbiology and Genetics, Georg-August-University Göttingen, Göttingen, Germany; Department of Biology, Chemistry, and Pharmacy, Institute of Chemistry and Biochemistry, Freie Universität Berlin, 14195, Berlin, Germany

**Keywords:** rAAV, AAV, digital PCR, genome quantification, genome integrity, heat-induced lysis artefacts, serotype differences

## Abstract

Adeno-associated virus (AAV) vectors are central to gene therapy, making precise genome quantification essential for product quality and dose determination. We systematically assessed digital PCR (dPCR) on the QIAcuity platform to quantify recombinant AAV2 and AAV8 genomes, examining how assay design, amplicon length, and heat-based sample preparation affect results. Across multiple genomic targets, shorter amplicons consistently yielded higher copy numbers than longer ones leading to a quantification deviation of up to ∼48%. The results point to heat-induced genomic fragmentation as the main cause of the observed effect. These findings highlight that dPCR-based AAV quantification is highly sensitive to amplicon length and sample preparation. We propose the use of the employed multitarget assay set to evaluate AAV genome extraction procedures, the quality of AAV genomic DNA extracts, and ultimately the AAV genome integrity.

## Introduction

Adeno-associated viruses (AAVs) are small, non-enveloped viruses that belong to the *Parvoviridae* family. AAVs contain a single-stranded DNA genome encapsulated within an icosahedral protein shell called a capsid. The genome is flanked by inverted terminal repeats (ITRs), which are crucial for the virus’s replication and integration into the host genome. To undergo a productive infection cycle, AAVs require co-infection with a helper virus, such as an adenovirus or herpesvirus. Without a helper virus, AAVs can integrate into the host genome, remaining latent until a helper virus activates them [1].

Unlike other viruses, AAVs are not known to cause disease in humans, making them a subject of intense research for therapeutic applications. To serve as a gene vector, the natural gene content of an AAV is replaced by a therapeutic gene flanked by the ITRs, resulting in a recombinant AAV (rAAV). The ability of rAAVs to infect both dividing and non-dividing cells and establish long-term expression of the delivered genes has positioned AAVs as one of the most promising vectors for gene therapy. Additionally, AAVs can target a wide range of tissues, including the eye, brain, liver, and muscle, allowing for the treatment of various genetic disorders. The long-term persistence of AAV-delivered genes is particularly beneficial for chronic conditions where sustained therapeutic expression is required [2–4].

Quantifying AAVs is a critical step in the development and manufacturing of AAV-based gene therapies, and it ensures the correct dosage is administered to patients, which is essential for safety and efficacy. Quantification typically involves measuring both the physical particles and the viral genomes. Physical particle quantification can be performed using techniques such as ELISA, which measures the capsid proteins, or nanoparticle tracking analysis, which counts the number of viral particles. Quantifying viral genomes is crucial to determine the number of functional viruses capable of delivering the therapeutic gene [5]. This is commonly done using quantitative PCR (qPCR), which amplifies and quantifies specific regions of the AAV genome. Digital PCR (dPCR) is another method that provides absolute quantification of viral genomes by partitioning the sample into thousands of partitions and performing PCR in each individual one [6]. Compared to qPCR, dPCR can provide absolute quantification and does not require a calibration curve, making it the method of choice [7]. However, the dPCR outcome significantly depends on the way how AAV samples are prepared and which assays are used [8].

The aim of this study was to evaluate a sample preparation procedure in combination with a dPCR assay, addressing several diverse regions of the rAAV genome for genome quantification and integrity assessment. While conducting our research, we were able to understand why AAV genome quantification depends on the employed assay, how this is related to sample preparation, and how the assays evaluated in this study can be used to evaluate sample preparation and data quality.

## Materials and Methods

### Digital PCR

Unless otherwise stated, all experiments were performed on the QIAcuity® dPCR system (QIAGEN) employing the QIAcuity® Probe PCR Master Mix (QIAGEN, 250101-250103) and QIAcuity® Nanoplate 8.5k 96-Well (Cat. No. / ID: 250021) for the gradient experiments and QIAcuity® Nanoplate 26k 24-Well (Cat. No. / ID: 250001) for other experiments. Raw data output was analysed with the QIAcuity® Software Suite (V2.5.0.0).

The temperature gradient was 64-52°C, and the cycling program included 2 minutes of initial denaturation at 95°C, up to 50 cycles with 15 seconds of denaturation at 95°C, and 1 minute of combined annealing and elongation. Imaging was realised after 40, 45, and 50 cycles with 500 ms exposure, and a gain setting of 6, for all channels.

Data analysis and visualisation were done using MS Excel or GraphPad Prism. For simplicity, the data are presented as genome copies per microliters as obtained from the dPCR reaction and are not converted to the estimated dosage units vector genome per millilitre.

### Used AAV material

Recombinant AAV8 and AAV2 were produced in HEK293 cells [9]. In the case of AAV8, the lysate was spiked with Denarase® (c-LEcta, Leipzig, Germany) at a final concentration of 50 U/mL. In the case of AAV2, the lysate was spiked with Benzonase® (Merck Milliporesigma, REF: E1014) at a final concentration of 25 U/mL. Both lysates were cleared via diatomaceous earth body feed filtration (C300) with a 0.22 µm PES filter [9]. The viral genome of AAV8 was 4261 bp, including both ITRs. The viral genome of AAV2 was 1944 bp, including both ITRs. Both genomes contained DNA fragments coding for a CMV promoter, CMV enhancer, and eGFP. For both AAV samples, as set, aliquots with 20 µL were prepared in 1.5 mL reaction tubes (Eppendorf, LoBind Tube, cat. No: 022431021) and frozen at -80°C until further use. A fresh aliquot was thawed for each experiment. Further, a full standard from Progen (AAV2 standard material (eGFP); Cat. No. 66V021; Lot No. FAK23070-01) and an empty standard from Progen (AAV2 standard material (eGFP); Cat. No. 66V020; Lot No. FAK23175-03) were used.

### AAV heat lysis

AAV heat lysis was done as described by Meierrieks et al. 2023 [8]. In brief, thawed at room temperature and diluted with the dPCR dilution buffer in a 1:10 manner. The appropriate dilution was incubated for 30 minutes at 95°C for AAV genome release.

### Primers and Probes

The primers and Probes used in this study are listed in Table 1 and were synthesised by Microsynth AG, Balgach, Switzerland. Probes were purified with HPLC, and primers were desalted after synthesis. Primer concentrations in the reaction mix were 0.8 µM and probe concentration was 0.25 µM.

**Table 1.**
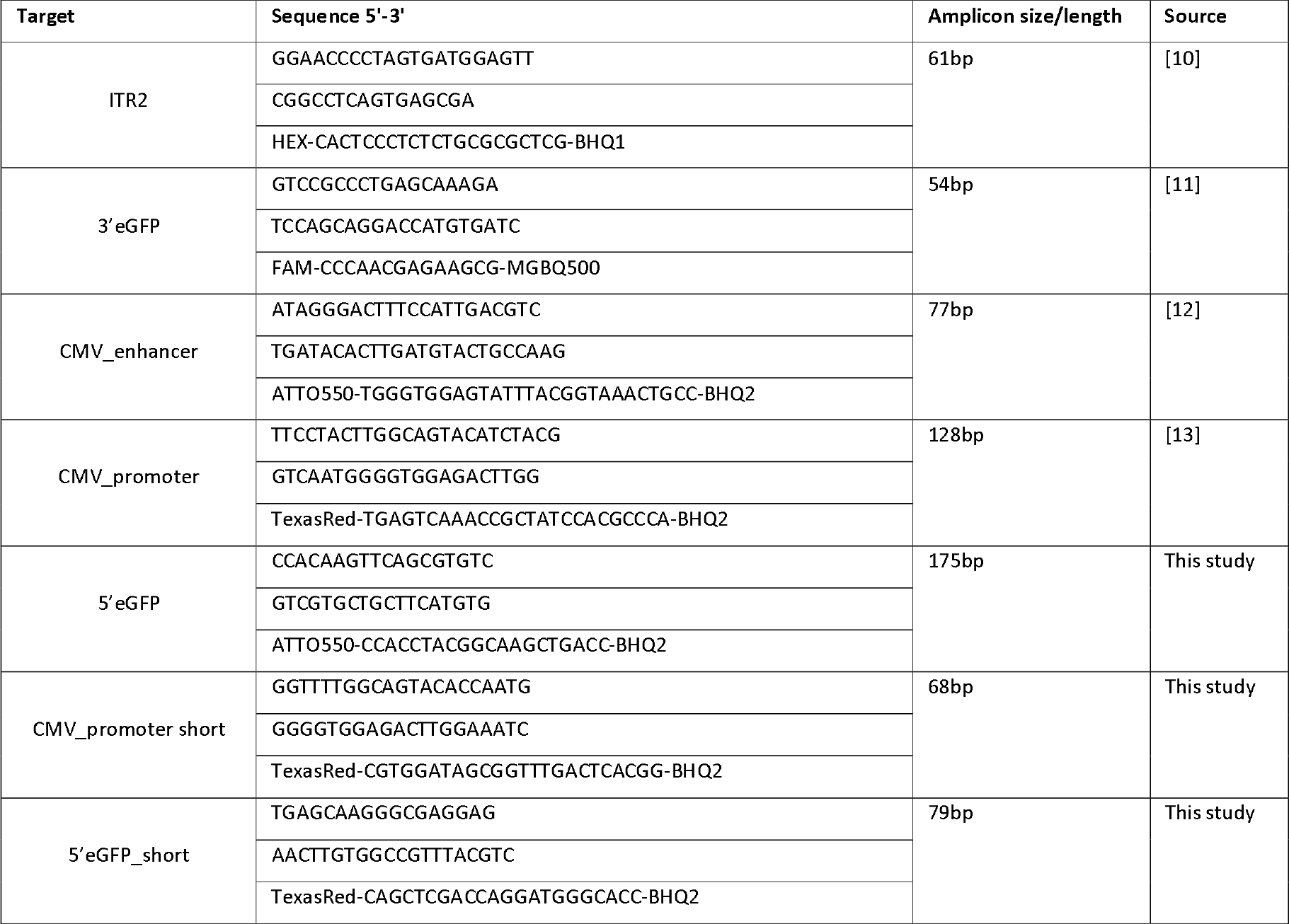
Primers and Probes.

## Results

### AAV Assay Evaluation

To reliably quantify the AAV genome and assess its integrity, all assays used must be evenly distributed across the genome (Figure 1 A) and, most importantly, perform consistently under the same experimental conditions. To identify the most suitable PCR parameters, we ran a gradient dPCR experiment employing all assays in combination with AAV2 and AAV8 samples as templates. We successfully identified universal PCR parameters for all assays. We assessed the cycling number by imaging after 40, 45, and 50 dPCR cycles. The one-dimensional scatter plot revealed a modest enhancement in signal separation for certain individual assays with increased cycling numbers, particularly evident for the ITR assay (Figure 1 B). However, after 40 cycles, signal separation was sufficient, and no further quantification improvement was observed with additional cycling. Consequently, 40 cycles at 57°C with a combined annealing and elongation time of one minute, lead to reliable results.

**Figure 1.**
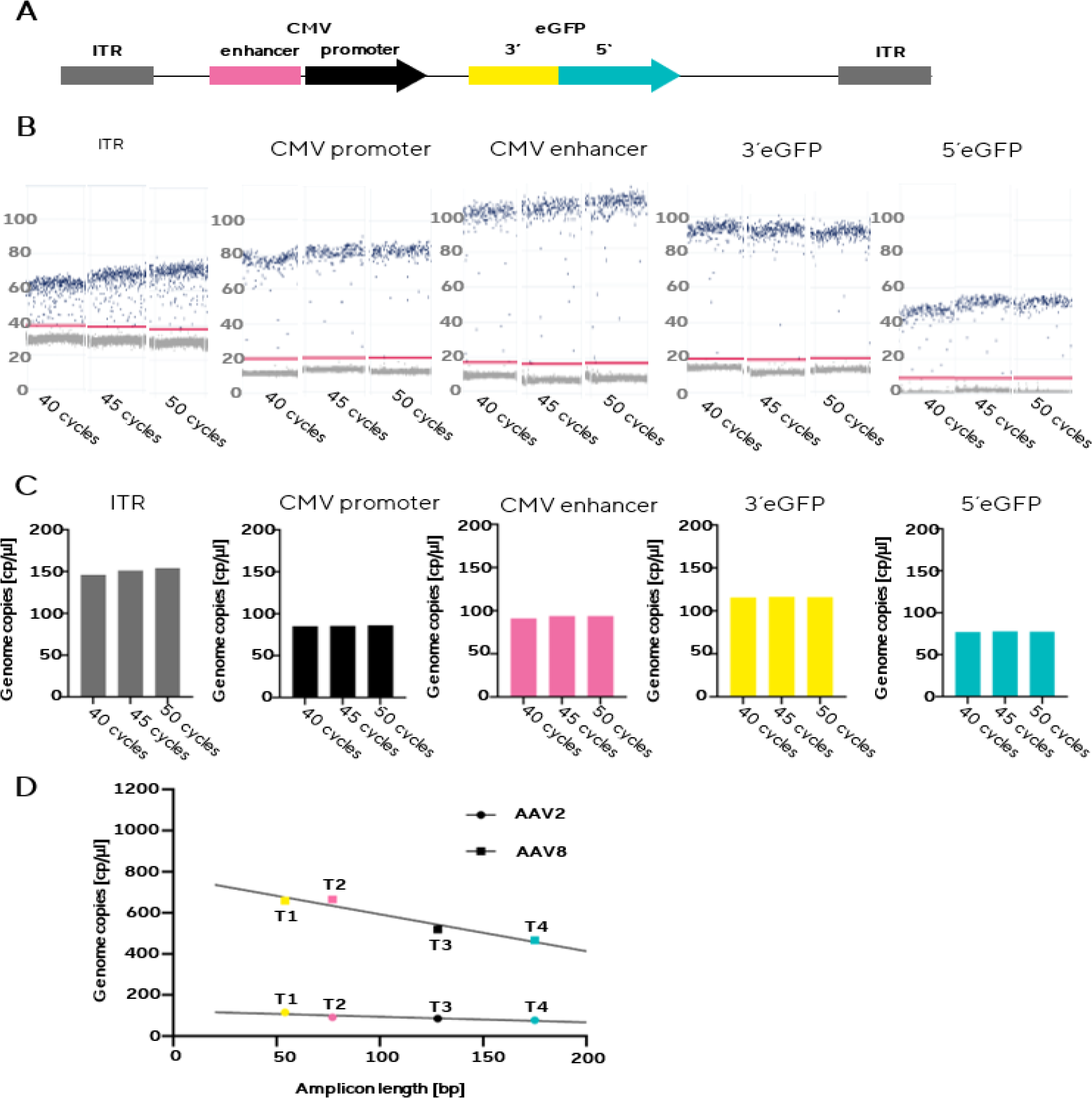
**A**: Distribution of used assays over the AAV genomes. The ITR regions flank both ends of the genome. **B**: One-dimensional scatter plot of 40, 45, and 50 cycles at 57°C. No relevant increase in fluorescence intensity with increased cycle numbers was observed. **C**: Genome copy numbers with 40, 45, and 50 cycles at 57°C. Determined assay specific genome counts at increased cycling numbers remained stable. However, assay-specific genome counts deviated. **D**: Correlation of determined genome counts plotted against the amplicon length of an assay. T1 = 3’eGFP, T2 = CMV enhancer, T3 = CMV promoter, and T4 = 5’eGFP. Both AAV2 and AAV8 serotypes faced heat lysis. A correlation between amplicon length and detected genome copies is obvious with both serotypes. The extent of correlations is serotype specific.

A notable observation from the initial experiments was the markedly diverse quantification of the utilized assays, despite employing the identical template input. The over-qualification of the ITR assay is well known [14]. Therefore, the data for this assay were omitted from the subsequent figures. However, the quantification discrepancy of approximately 50% between the 5’eGFP assay and the 3’eGFP assay, both targeting the same transgene, was unexpected. The other assays employed also yielded distinct quantifications, none of which matched were comparable (Figure 1 C). When correlating the amplicon length with the determined genome counts, the link between quantification of AAV genomes and amplicon size becomes evident. To be more precise, the longer the amplified fragment, the lower the quantification, and vice versa, the shorter the amplified fragment, the higher the quantification. This phenomenon was confirmed in both AAV2 and AAV8 samples (Figure 1 D), demonstrating its independence from serotype or genome.

These findings point out the likelihood of no AAV assay quantifying the true genome count, as their precision depends on the amplicon size, which in turn is dependent on the given sequence composition of the target itself.

### The underlying root cause

Genome counts of AAV2 and AAV8 revealed a linear correlation with the amplicon size, but with a different slope (Figure 1 D), implying the underlying root cause for the observed effect. It was reported that different AAV serotypes exhibit different levels of heat resistance [15]. According to Bennett and colleagues [15], the AAV2 virion is more resistant to heat compared to the AAV8 virion in a tris-based buffer system. Such a buffer system was also employed in these experiments as dilution buffer and during virion lysis. Consequently, AAV2 releases its genome later during the heat lysis process compared to AAV8. This leads to the hypothesis that heat impacts the viral genome during virion lysis. Assuming a random shearing, larger genomic target regions would be affected more frequently leading to artificially reduced quantification if compared to shorter genomic target regions.

To challenge this hypothesis, two additional assays were designed, targeting the same regions as the 5’eGFP assay and the CMV promoter assay, but resulting in a shorter amplicon. Thus, if the quantification differences truly reflect the degree of shearing of the viral genome rather than the sequence composition of the amplified region, the obtained viral genome copy numbers should align with the previously observed linear correlations. Results perfectly matched the AAV genome quantification / amplicon size correlation and confirmed the presented hypothesis (Figure 2 A). Both assays with shorter amplicons yielded higher genomic titers compared to their longer counterparts. Specifically, the CMV promoter short amplicon version showed a ∼19% increase in genome count compared to the CMV promoter long amplicon version. Likewise, the 5’eGFP short amplicon assay exhibited a ∼48% increase in genome count compared to the 5’eGFP long amplicon assay. The 3’eGFP assay with the shortest amplicon was used as reference.

**Figure 2.**
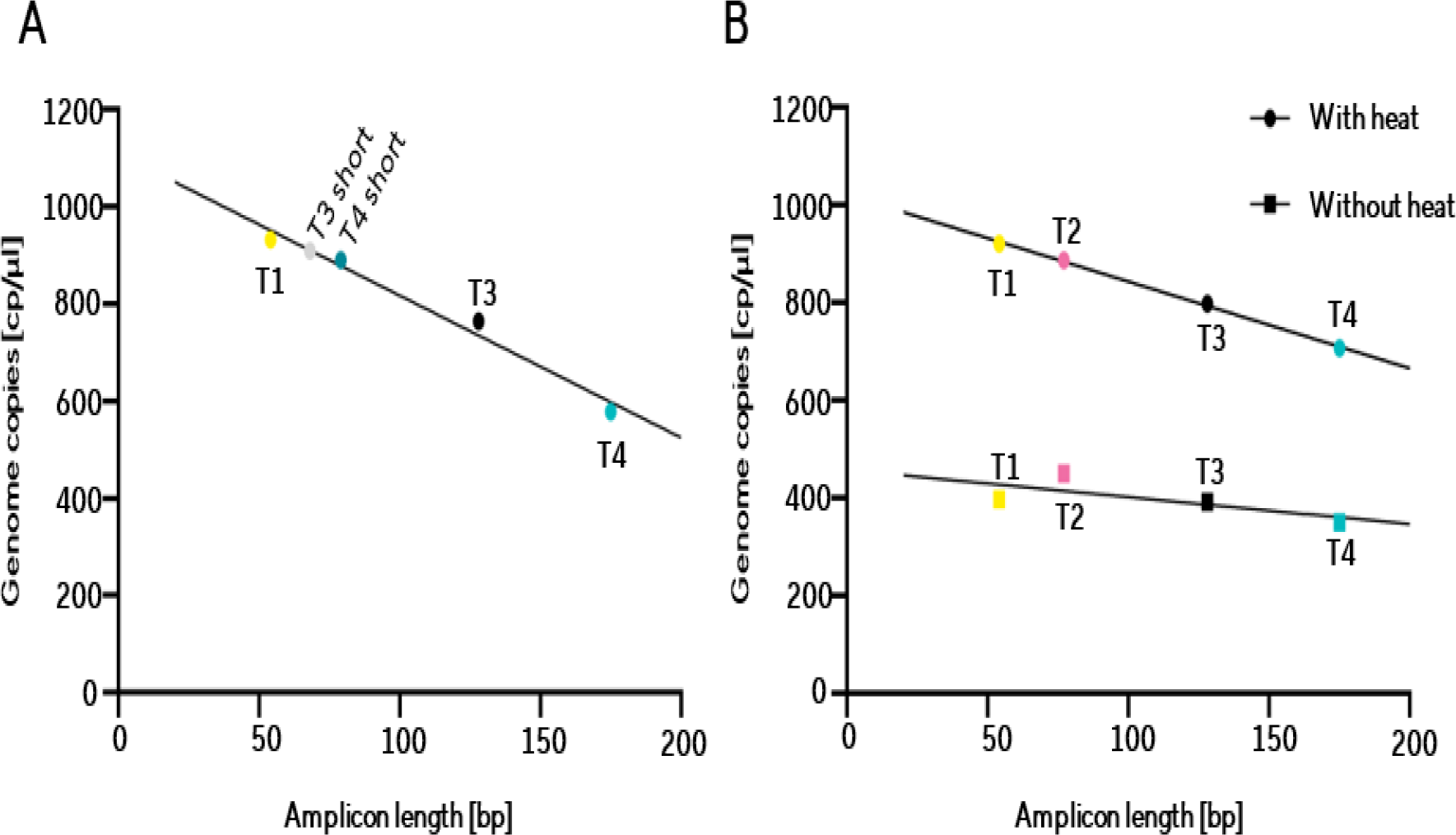
**A**: Correlation of determined genome counts plotted against the amplicon length of an assay. T1 = 3’eGFP, T2 = CMV enhancer, T3 = CMV promoter, T4 = 5’eGFP, including an assay targeting the CMV promoter region but leading to a shorter amplicon and an assay targeting the 5’eGFP region but leading to a shorter amplicon. **B**: Correlation of determined genome counts against the amplicon length of an assay on a heat-treated sample and on a native sample.

To challenge the counterhypothesis “*Absence of heat reduces deviations between the different assays*”, we quantified natively AAV8 virions using dPCR without prior heat treatment (Figure 2 B). Results revealed a reduced genome count overall, which likely originates from aggregated viral particles [16,17]. However, this should not affect the conclusion of the experiment in terms of the heat effect, where a reduced spread of the obtained genome counts with different assays was observed.

In summary, the results prove that heat-induced viral genomic DNA damage is the primary cause of the observed quantification deviations. Thus, the meticulously curated set of assays functions as a potent instrument for evaluating the quality of input AAV DNA.

## Discussion

The key finding of this study is that heat is highly relevant for sample preparation of AAV for dPCR-based genome quantification. Although DNA is very stable under physiological conditions [18], this series of experiments showed that ssDNA AAV genomes are non-specifically fragmented by heat during sample preparation, meaning that they can no longer be quantified accurately. The anticipated precision of digital PCR is compromised by heat-associated sample preparation in combination with an AAV assay that utilizes a larger amplicon. Heat-induced genome degeneration not only affects quantification but also the integrity assessment of AAV preparations. The addition of nuclease in the AAV process to remove resDNA can, of course, also be suspected as the cause of the detected phenomena. We rule this out due to the fact of not observing any degradation of the probe during the measurement of native virions (Figure 2 B). In addition, purified AAV standard material was analyzed by an independent group using the same assay set and heat lysis procedure. Results show the same dependence on the amplicon size with a comparable spread of the determined genome numbers.

Heat-induced DNA degradation is a complex process involving many factors. The two most relevant to PCR are the hydrolysis of phosphodiester bonds, which directly leads to the physical disruption of the DNA strand, and the hydrolysis of glycosyl bonds, which decouple the bases from the ribose, thereby limiting primer recognition of the template and PCR processivity [19]. Other factors contributing to heat-induced degradation include heat-induced deamination of the bases [20,21], which promotes the formation of nitrogen oxides [22] and ROS [23]. Although these effects mainly occur in acidic and neutral environments, they have also been observed in basic TE buffer, similar to the one used here.

As a direct logical consequence, the heat must be removed from the process. In our procedure, the capsids are denatured with heat to release the viral genomes. This can also be done enzymatically using proteinase K, but here too, the proteinase K would have to be inactivated again using heat. Additionally, heat cannot be completely removed from the system, since in PCR, an initial denaturation step at 95°C is necessary to activate the hot start polymerase. So, how can the amount of heat in the system be reduced? The use of chaotropic salts, as described by Boom and colleagues [24], is a promising approach. The lysis buffer contains guanidine thiocyanate and a PCR-compatible surfactant. It does not contain any antioxidants to counteract heat-induced oxidative stress. However, a further development of this lysis buffer by Kravchenko and colleagues [25] also includes 2-mercaptoethanol as a reducing agent. The chaotropic lysis buffer allows DNA isolation from a whole blood sample without heat treatment and the preservation of the nucleic acid for over a year at - 20°C. Therefore, lysing AAVs with a chaotropic lysis buffer may circumvent heatlZlinduced issues. Whether this can be done entirely without heat must be verified empirically in further studies.

The assay set established in this investigation can also be used in multiplex without restriction. It is an ideal tool for evaluating potential novel approaches to AAV lysis without or with reduced heat exposure. Until further advancements are made, it is conceivable to employ the assay set to estimate the actual genome titer by extrapolating the regression line. The variation in the slope across different experiments allows to evaluate reproducibility. Genome integrity can also be estimated with the assay set, once through the consistency of the quantification across all assays and through a co-localization analysis [26].

## Acknowledgement

Many thanks to the Lab Separations Applications team for the Lab Essentials Applications Development department - Campus Göttingen for providing AAV samples.

## Authors contributions

RH designed the study; LK, MG, RH designed assays; AG, FT, NW performed experiments; AG, RH analysed data and wrote the manuscript; AMS and KP ensured funds.

## Declaration of Generative AI and AI-assisted technologies in the writing process

The authors used DeepL Translate, DeepL Write, and Sartorius GPT-4 Turbo Chat AI to improve language and readability. AI output was reviewed by the authors before including in the manuscript. The authors take full responsibility for the content of the publication.

